# Swimming kinematics and performance of spinal transected lampreys with different levels of axon regeneration

**DOI:** 10.1101/2021.03.26.437228

**Authors:** Jacob. Fies, Brad J. Gemmell, Stephanie M. Fogerson, John H. Costello, Jennifer R. Morgan, Eric D. Tytell, S. P. Colin

**Author notes:** Summary statement: We show that lampreys who have recovered from having their spinal cords transected do not fully regain swimming abilities are not able to swim as efficiently as non-transected lampreys.

## Abstract

Neural and functional recovery in lampreys from spinal cord transection has been well documented. However, the extent of axon regeneration is highly variable and it is not known whether it is related to the level of behavioral recovery. To address this, we examined how swimming kinematics were related to axon regeneration by quantifying the relationship between swimming performance and percent axon regeneration of transected lampreys after 11 weeks of recovery. We found that swimming speed was not related to percent axon regeneration but it was closely related to body wave frequency and speed. However, wave frequency and speed varied greatly within individuals which resulted in swimming speed also varying within individuals. In fact, most recovered individuals, regardless of percent axon regeneration, could swim at fast and slow speeds. However, none of the transected individuals were able to generate body waves as large as the control lampreys. In order to swim faster, transected lampreys increased their wave frequencies and, as a result, transected lampreys had much higher frequencies than control lamprey at comparable swimming velocities. These data suggest that the control lampreys swam more efficiently than transected lampreys. In conclusion, there appears to be a minimal recovery threshold in terms of percent axon regeneration required for lampreys to be capable of swimming, however, there also seems to be a limit to how much they can behaviorally recover.

## Introduction

Anguilliform propulsion has been shown to be one of the most efficient forms of swimming propulsion observed among animals (Van Ginneken et al., 2005). This form of propulsion is characterized by a traveling wave that moves from the head to the tail with a relatively short wavelength, so that about a full wave is present on the body at any time. The amplitude of this traveling wave increases as it travels down the body (Lauder and Tytell, 2005). These kinematics interact with the adjacent fluid to slowly build fluid vorticity and strong negative pressure regions that serve to efficiently generate a suction thrust that pulls the anguilliform swimmer forward (Gemmell et al., 2016).

Larval sea lampreys (*Petromyzon marinus*) are well characterized anguilliform swimmers (McClellan et al., 2016). Healthy lampreys generate the characteristic traveling wave using muscle contractions along the side of their body that are initiated just caudal to the head and travel toward the tail. By alternating these contractions on each side of the body the lamprey can generate successive traveling waves that make up each swimming cycle (McClellan, 1989; Williams, 1989; Williams and McMillen, 2015). The speed of the observed body wave is slower than that of the muscle contraction as a result of the interaction of forces acting on the body which include the forces generated by the muscles and the resistive forces of the fluid acting on the body (Ding et al., 2012; Tytell et al., 2010; Williams, 1989; Williams and McMillen, 2015) Demonstrating the robustness of this behavior, within several months after a complete spinal cord transection, lampreys are able to achieve robust recovery of swimming behaviors (Cohen et al., 1999; Hanslik et al., 2018; Katz et al., 2020; McClellan, 1989; McClellan, 1994; Oliphint et al., 2010; Rovainen, 1976; Selzer, 1978). Remarkably, they can also recover normal swimming after a second spinal re-transection (Hanslik et al., 2018). Therefore, lampreys have served as a great model for studying anguilliform swimming, as well as recovery from spinal cord injuries. Lampreys spontaneously recover swimming behaviors within 8-12 weeks after their spinal cord is transected rostrally at the level of the 5th gill due, in part, to long-distance regeneration of descending axons (McClellan, 1994; Oliphint et al., 2010; Rovainen, 1976). Initially, such transected animals are completely paralyzed (Hanslik et al., 2018; Oliphint et al., 2010). Axon regeneration begins two to three weeks after spinal cord transection with axons beginning to regenerate and some observed locomotor function just caudal to where the spinal cord was transected. Progressively over time, locomotor function can be observed more caudally and by 8-12 weeks locomotor activity, and neural activity and movement patterns can be similar to normal, healthy larval lamprey (Cohen et al., 1986; McClellan, 1994; Oliphint et al., 2010). Despite the ability to regenerate axons, the swimming kinematics and performance of recovered lampreys still differs from non-transected lampreys (Oliphint et al., 2010). In addition, only 30-70% of descending reticulospinal (RS) axons regenerate, making a few, sparse synaptic connections, implicating compensatory mechanisms in locomotor recovery (Davis Jr and McClellan, 1994; Oliphint et al., 2010; Shu Yin and Selzer, 1983).

Lampreys that have more caudal spinal cord transections (mid-body or lower) can often swim immediately after transection. Under these conditions, locomotor activity is only present rostral to the lesion, and the body waves are passively propagated to the caudal region (Gemmell et al., 2016; McClellan, 1990). In comparison to rostral lesions, axon regeneration was reported to be less robust after transection of the caudal spinal cord (Shu Yin and Selzer, 1983).

Despite lampreys being well documented as robust regenerators after a rostral spinal cord transection (e.g. in the gill region) and progressing from paralysis to full mobility within 10-12 weeks post injury, the extent of axon regeneration supporting this behavioral recovery is variable from animal-to-animal (Cohen et al., 1986; Hanslik et al., 2018; Oliphint et al., 2010; Rovainen, 1976; Selzer, 1978). Therefore, to understand the relationship between neural regeneration and behavioral recovery, we quantified the kinematics and swimming abilities of larval lampreys that had different levels of axon regeneration across the site of spinal transection and compared performance of control lamprey (non-transected) to lamprey who recovered from having their spinal cord transected 10.5 weeks prior at the 5^th^ gill or at mid-body.

## Methods

### Spinal cord transections

All animals used in the experiments were late larval-stage lampreys (*Petromyzon marinus*; 10–14 cm; M and F) that were housed at room temperature (25 °C) in 10-gallon aquariums. Fourteen lamprey (treatments) underwent spinal cord transection surgery (n= 11 were transected at the 5^th^ gill and n= 3 at midbody) as previously described by Oliphint et al. (2010). Briefly, each lamprey was anesthetized with Finquel MS-222 (0.1 g/L tank water; Argent Chemical Laboratories) and then placed in a Sylgard-lined Petri dish on a paper towel moistened in oxygenated lamprey Ringers (100 NaCl, 2.1 KCl, 2.6 CaCl_2_, 1.8 MgCl_2_, 4 glucose, 0.5 glutamine, 2 HEPES, pH 7.4). A dorsal incision was made either at the 5^th^ gill or approximately halfway down the length of the body, just above the dorsal fin, through the skin, musculature and fat tissue in order to expose the spinal cord. Then, the spinal cord was completely transected either at the 5^th^ gill or the mid-body with a single horizontal cut made with fine iridectomy scissors (Fig. 1A). The incision was closed with a single suture (Ethilon 6-0 black monofilament nylon; Johnson & Johnson, Langhorn, PA). Three control animals received a sham treatment in which they underwent the same surgical procedures, but the spinal cord was not cut (Table 1). All animals were then returned to their home tanks for 11 weeks post-injury until they were video recorded. All procedures were approved by the Institutional Animal Care and Use Committee at the Marine Biological Laboratory in accordance with the standards set by the National Institutes of Health.

**Fig. 1.**
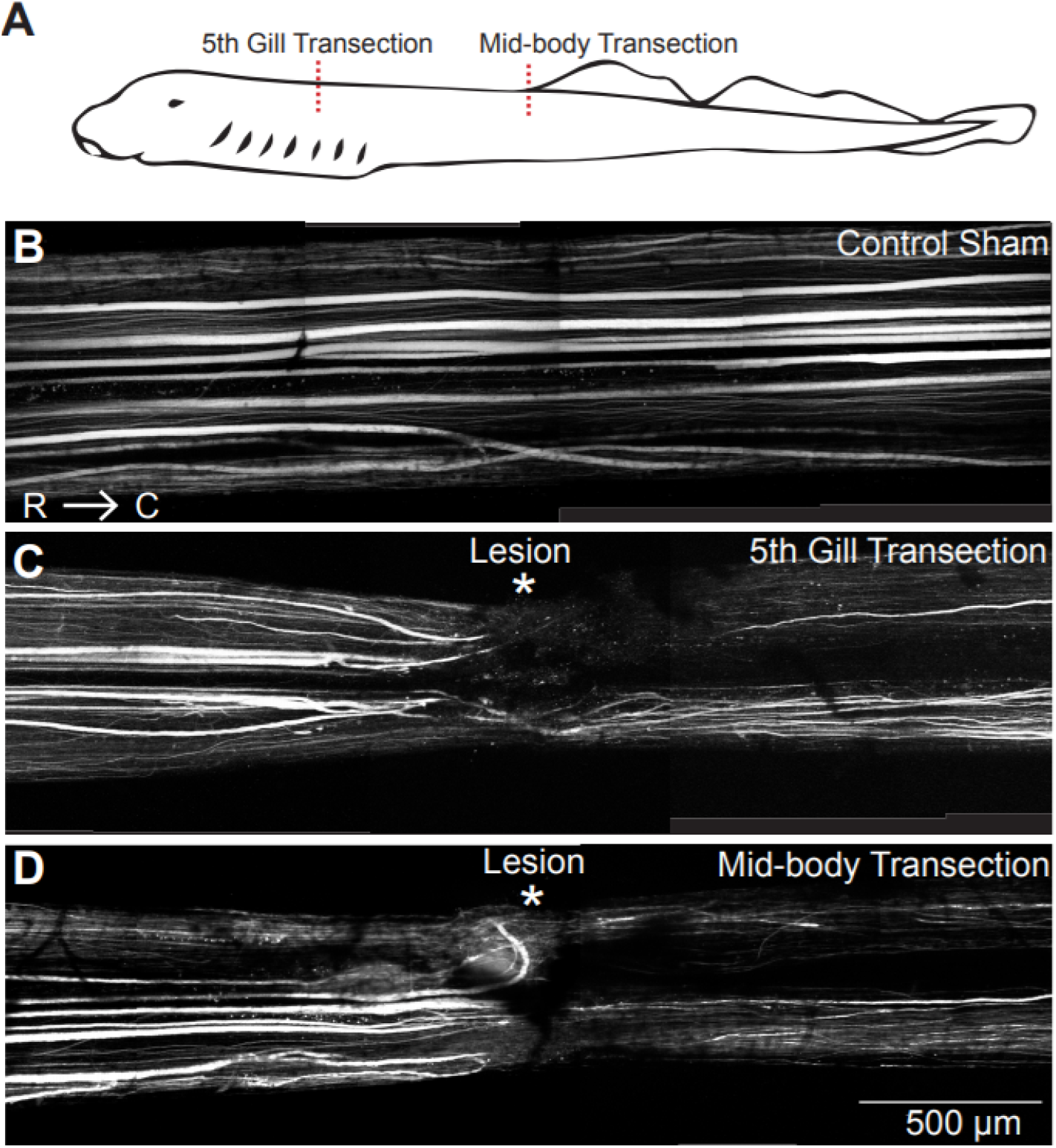
Bulk labeling of regenerating axons ~10.5 weeks following a full spinal cord transection. A) Schematic of a larval lamprey with the site of spinal cord transection indicated with a red dashed line for a 5th gill or mid-body transection. B) A montage of confocal z-projections stitched together of a control, sham uninjured spinal cord with axons labeled by a 10kaDa Alexa Fluor 488 dextran, showing fairly uniform labeling along the length of the spinal cord. C-D) Labeling of axons ~ 10.5 weeks post injury in a 5th gill transected and a mid-body transected animal shows sparser axon labeling in the region caudal to the lesion site in comparison to the rostral region, indicating the amount of axon regeneration. Note that the amount of axon regeneration is comparable between the 5th gill and mid-body transected spinal cords. Scale bar in D applies to panels B-D. Rostral (R) is to the left and caudal (C) is to the right.

### Video recording and Kinematics Calculations

At 11 weeks post-injury, videos were taken of the lampreys as they were prompted to swim through a 1.5 × 5 m acrylic aquarium that was filled with 5 cm of lamprey tank water. Video was captured at 1000 fps using a Photron Fastcam 1024 PCI video camera positioned below the lampreys (as in Gemmell *et al*., 2016).

To compare the kinematics and swimming of the lamprey, each animal was video recorded during steady state swimming according to Gemmell et al. (2016) and (Du Clos et al., 2019)Du Clos et al. (2020). Accordingly, lampreys were placed at one end of long (1.5 meter) tanks where swimming was initiated by touching the individual gently at the tail. Swimming and kinematics was videoed as the lamprey passed the middle of the tank (no longer accelerating and swimming in steady state). Their bodies were illuminated with a light sheet that was oriented horizontally and directed perpendicular to the camera angle, and was generated using two lasers (532 nm, 600 mW continuous wave per laser) placed on opposite sides of the aquarium. Using two lasers eliminated shaded regions around the swimming lampreys and enabled us to thoroughly illuminate the outline of the lamprey. The laser light did not affect the lampreys’ swimming behaviors as the animals were larvae and thus still retained tissue covering their eyes. Only video sequences where the velocity averaged over the entire sequence remained constant were used in the analysis.

Swimming kinematics were quantified manually using ImageJ (NIH) software and an in-house MATLAB program (https://github.com/tytell/neuromech.wiki.git). Raw images of the freely swimming animals were input to a custom program in MATLAB that identified and tracked the midline of the lampreys as they swam. Based on the X, Y coordinates of the lamprey midline the max amplitude, wavelength, and frequency was calculated over time. Maximum amplitude was calculated at the highest point in the wave along the body. Wavelength was done similarly but between wave peaks or troughs.

To arrive at estimates of relative efficiency based on kinematics we calculated Strouhal number (St) as St = 2fA/U, where A is the maximum amplitude, f is the frequency and U is the swimming speed (Triantafyllou et al., 1993; Tytell, 2004). We also calculated the stride length, the distance traveled per body wave, by dividing swimming speed by wave frequency to get an estimate of how effective each body wave was at propelling the lamprey forward.

### Axon labeling, Imaging and Regeneration analysis

Following video recording at 10.5 weeks post injury, the descending reticulospinal axons were bulk labeled in order to assess the extent of axon regeneration, as previously described (Armstrong et al., 2003; Hanslik et al., 2018; Lau et al., 2013). Briefly, animals were re-anesthetized in MS-222, and a second spinal lesion was made 0.5 cm rostral to the original transection site. A 1×1×1 mm cube of Gelfoam (Pfizer; New York, NY) soaked in 5 mM Alexa Fluor® 488-conjugated dextran (10kDa; Thermo Fisher, Inc. Waltham, MA), diluted in lamprey internal solution (180 mM KCl, 10mM HEPES, pH7.4), was placed in the lesion which was then closed with a single suture. Spinal cords were harvested three days after labeling to allow for maximum transport of dye. The anterograde-labeled, regenerating axons were imaged live within whole mounted spinal cords submerged in oxygenated lamprey ringer. Imaging was performed using a Zeiss LSM 510 laser scanning confocal on an Axioskop 2FS upright microscope (10x, 0.3 NA Zeiss EC Plan-Neofluar objective). Z-stacks of spinal cords were acquired at distances ranging from 2 mm proximal to the original transection site to 5 mm distal. Maximum intensity projections were made using the Zeiss LSM software and stitched together in Photoshop. For fifth gill and mid-body transections, the number of labeled axons crossing fiducial markers positioned at 1-1.5 mm proximal and 1.0 mm distal to the transection site were counted. Percent axon regeneration was calculated as the number of labeled axons distal to the transection site, divided by the number of labeled axons at rostral, though we acknowledge that this is a semi-quantitative estimate that may include some axon branches. Controls spinal cords were imaged and analyzed the same way, except without an intervening lesion site.

### Statistics

All comparisons were tested to determine if they complied to the assumptions of parametric tests. Wave kinematics were compared among treatments (5^th^ gill transected, mid-body transected and sham) using One-way ANOVAs. The relationships between axon regeneration and swimming performance and kinematics were examined using regression analyses. We additionally examined swimming speed as a function of axon regeneration and tail beat frequency using a mixed model multiple regression, including regeneration, tail beat frequency, and their interaction as fixed factors and individual animal as a random effect. This statistical model was implemented using R 4.0.2 and nlme 3.1−148 (Pinheiro et al., 2007)

## Results

### Axon regeneration after spinal cord injury in lampreys

For this study, we examined and compared axon regeneration, kinematics, and performance in lampreys that underwent spinal cord transection at the level of the 5^th^ gill or the mid-body (Fig.1A). In untransected control spinal cords, RS axons generally projected in relatively straight patterns within the ventromedial and ventrolateral tracts (Fig. 1B). In contrast, at ~10.5 weeks post-injury, RS axons proximal to the spinal lesion can be straight, curved, or branching within the spinal cord (Oliphint et al., 2010), and only a subset of RS axons regenerated distal to the lesion (Fig. 1C). Similar amounts of RS axon regeneration were observed in spinal cords that were transected at the mid-body (Fig. 1D), a perturbation that does not result in paralysis of the animal due to preservation of the rostral spinal circuits that initiate swimming (Gemmell et al., 2016). To estimate the extent of axon regeneration, we counted the number of Alexa-Fluor® 488-labeled RS axons 1 mm distal to the lesion center and divided this by the number of labeled axons 1-1.5 mm proximal to the lesion. In this cohort of animals, the percent axon regeneration in spinal cords was between 33.3 and 84.2 percent with a median of 58.6 percent regeneration, which is similar to that reported in previous studies (Lau et al., 2013; Oliphint et al., 2010; Shu Yin and Selzer, 1983), thus providing a range of neural regeneration to compare to behavioral performance.

### Swimming performance and kinematics

All of the lampreys examined recovered sufficiently to be able to swim. In fact, we found that how fast the lampreys were capable of swimming was not related to the percent of axon regeneration within their spinal cords (Fig 2A; Regression analysis, df = 1, F = 2.02, P = 0.2). Comparison of swimming speeds among treatments (5^th^ gill transected, mid-body transected and sham control) suggests that the control sham lampreys swam faster than transected individuals but the differences were not significant (Fig. 2B; ANOVA, df = 2, F = 2.72, P = 0.09). A more quantitative comparison of body kinematic variables to percent regeneration shows that none of the wave kinematics were significantly related to axon regeneration for the recovered transected lampreys (Fig. 2C-E, Regression analysis, df = 1, P > 0.07). However, the control sham lampreys had significantly longer wavelengths and higher amplitudes than transected lampreys (Holm-Sidak post-hoc comparison, p < 0.05), but their wave frequencies were not different than transected lampreys (ANOVA, df = 2, p > 0.1).

**Fig. 2.**
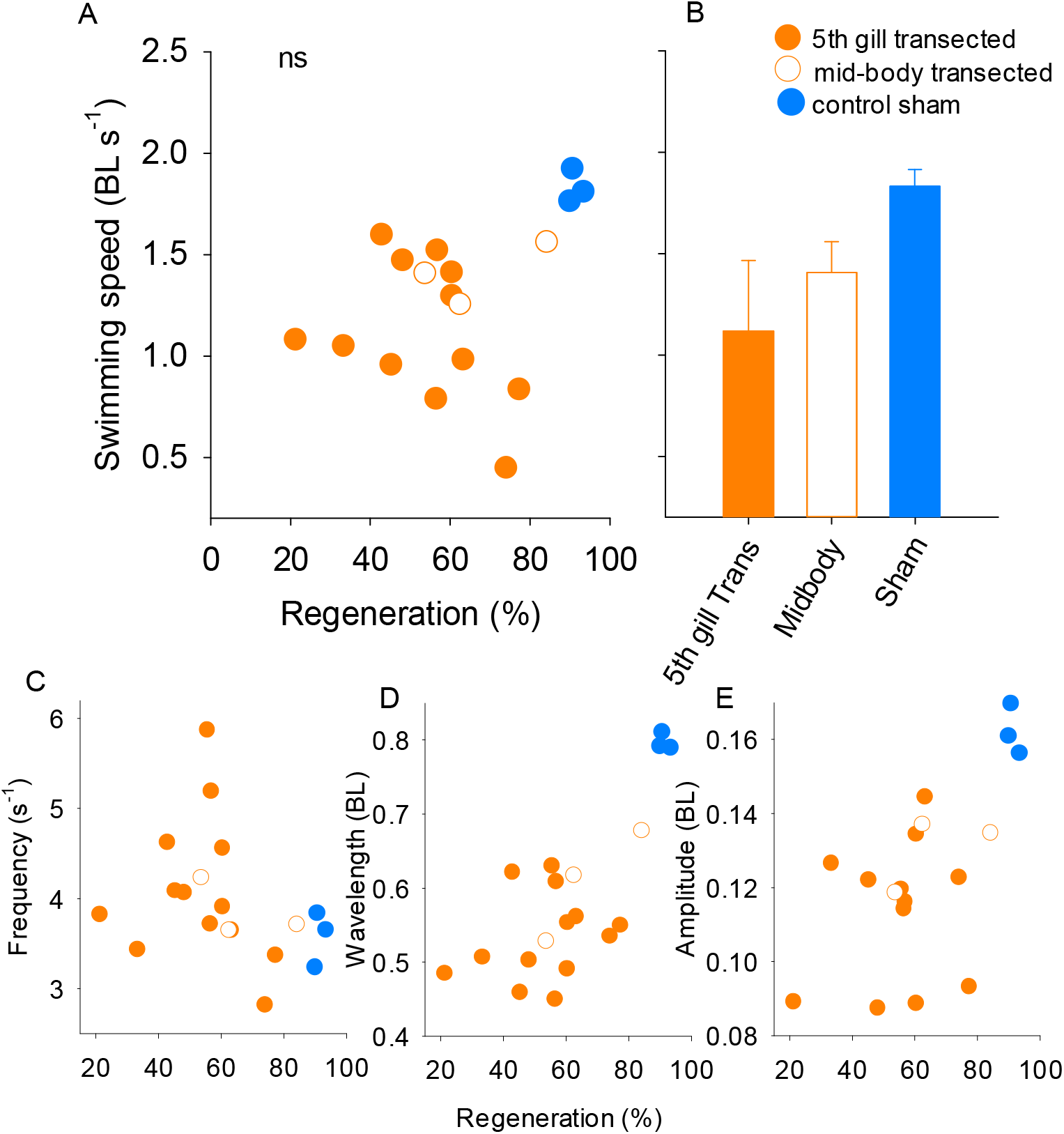
A) Swimming speed of lampreys versus their degree of spinal cord regeneration (%) after recovering for 10.5 weeks from having their spinal cord transected (Regression analysis, df = 1, p > 0.05). B) Comparison of mean swimming speeds among treatments (ANOVA, df = 2, p > 0.05). C-E) Comparison of A) wave frequency, B) wavelength and C) wave amplitude versus the degree of spinal cord regeneration (%; Regression including 5ht gill and midbody, df = 1, p > 0.05).

Despite the lack of correspondence between spinal cord regeneration and swimming performance, a visual comparison of representative recovered individuals to a control (or sham) lamprey illustrates that there were important differences in the body and swimming kinematics that swimming speed did not capture. Sequential images of the lampreys (Fig. 3A) reveals that the wavelength of the body wave of the control lamprey was large compared to the recovered, transected lampreys. As such, more waves occurred along the bodies of the transected lampreys (1.9 ± 0.2 waves per body) at any one time than the control lampreys (1.2 ± 0.01). The swimming kinematics of the control lamprey were also very regular during consecutive swimming cycles, while the swimming kinematics of the transected lampreys were much more irregular (seen in the motion of the head and swimming velocity (Fig. 3B and D)). A fast swimming 5^th^ gill transected individual was included in the comparison to illustrate the differences in the kinematics between the fast transected and the control (Fig. 3). Despite traveling a similar distance as the control (Fig. 3C), the transected lamprey still had a smaller wavelength (Fig. 3A), the head moved back and forth much more frequently (Fig. 3B) and the swim pattern was much more erratic than the control (Fig. 3D).

**Fig. 3.**
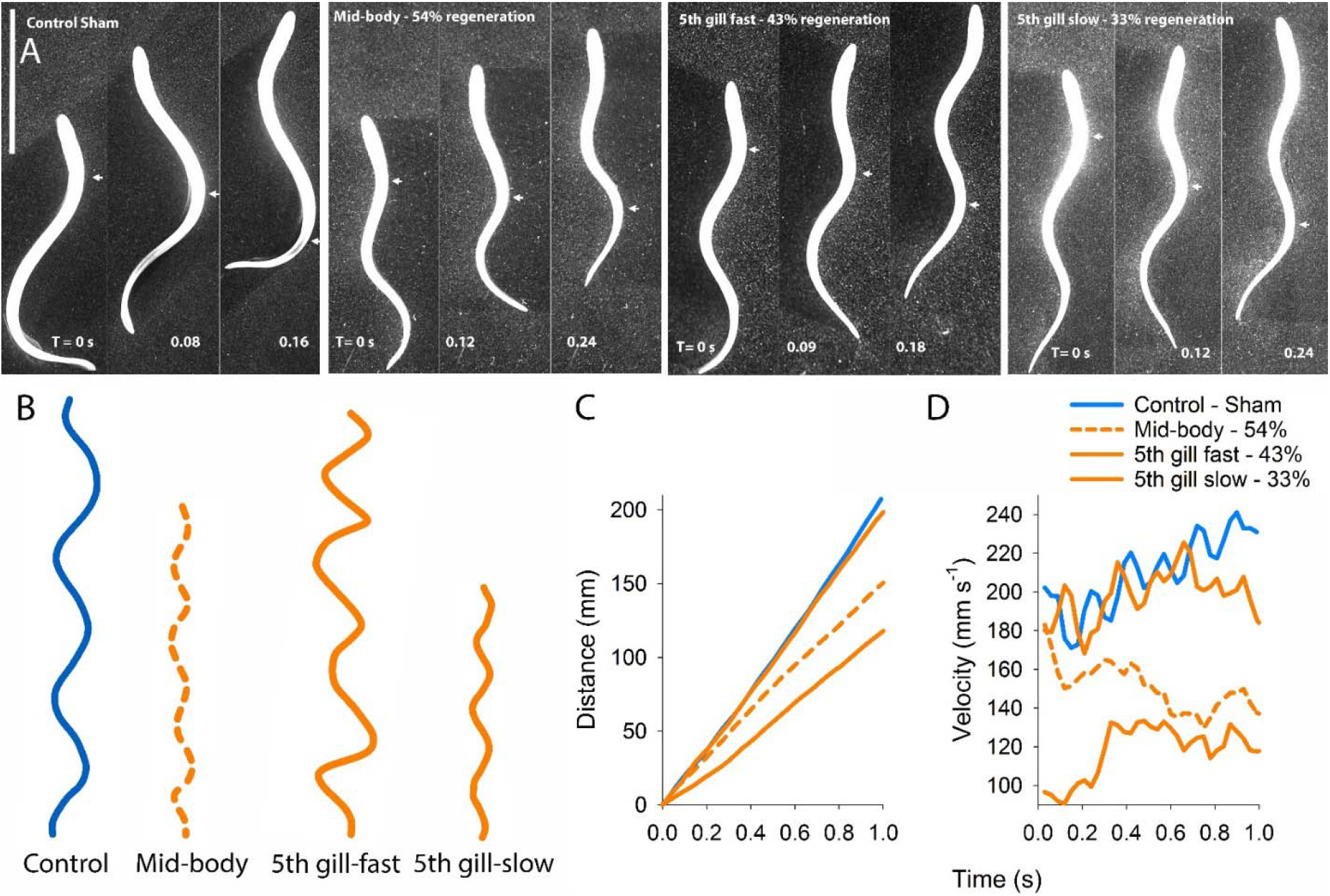
Comparison of body and swimming kinematics of lampreys. A) Sequential images of different lamprey showing the progression of a body wave (indicated by white arrow) moving from head to tail. Notice the control lamprey has only one large wave traveling along the body at a time while all the transected lampreys, regardless of swimming velocity (D), have multiple smaller waves moving along the body. B) Tracking of the movement of the head of the lamprey for 1 second. Notice the distance traveled and the evenness vs. uneveness of the lateral motion of the heads through time. C) Distance the different lamprey traveled over a second. D) Velocity of the different lamprey over a second. Notice the regular swim cycles of the control lamprey (blue) versus the more erratic motion of the lampreys transected (orange).

A closer look at the wave amplitudes among the groups revealed that the larger amplitudes observed for the control lampreys were achieved by the lamprey increasing amplitude as the wave traveled head to tail (Fig. 4). In contrast, the wave amplitudes of the 5^th^ gill and mid-body transected lampreys did not change as much as the waves moved head to tail.

**Fig. 4.**
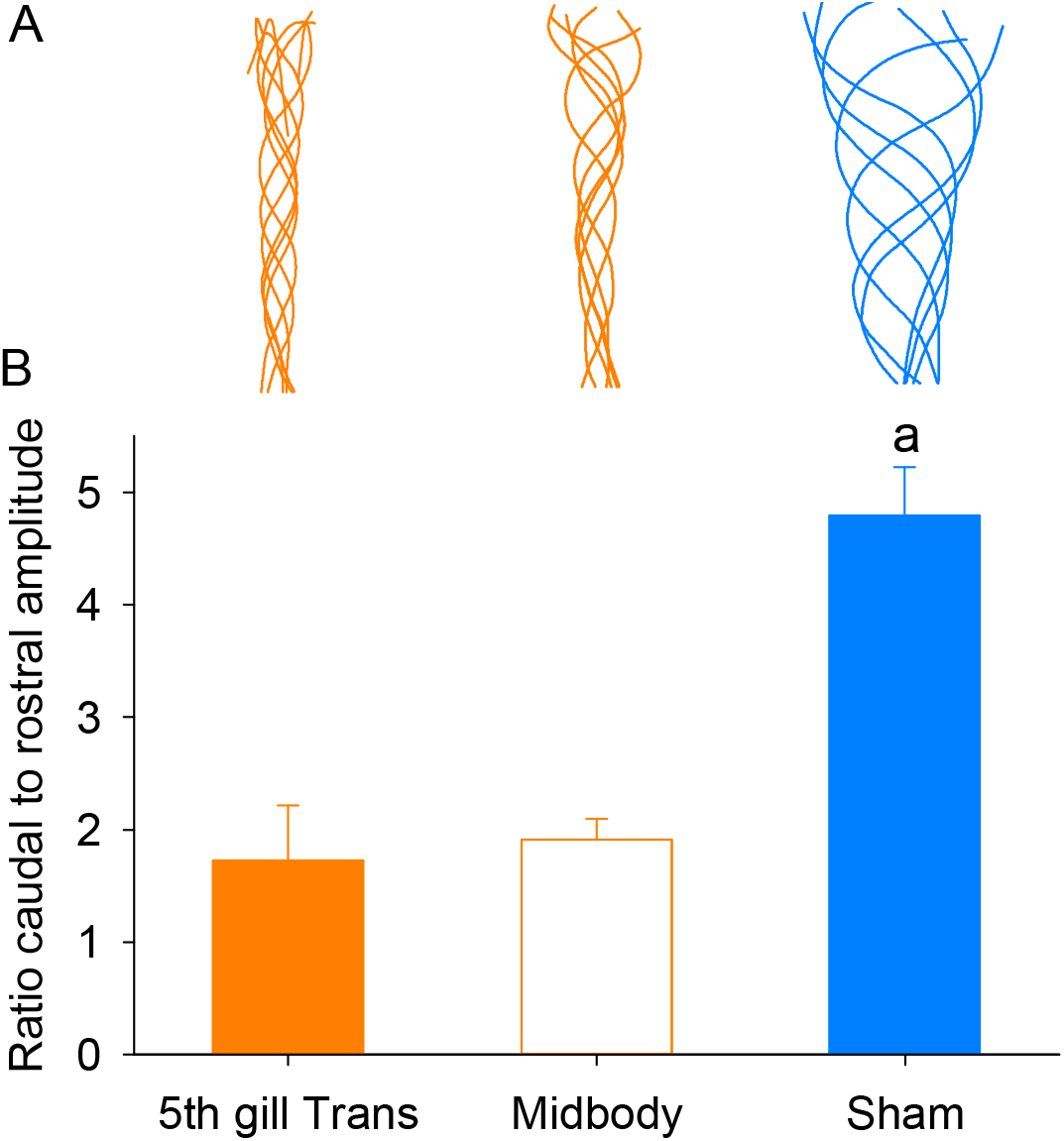
Change in body wave amplitude as wave travels from head to tail. A) Change in midline of representative lampreys over time. B) Mean change in amplitude (as ratio of amplitude at the tail (caudal) and the head (rostral)) among treatments. Lower case letters significantly different treatment groups (Holm-Sidak post-hoc comparison, p < 0.05).

While average swimming speed was not significantly related to axon regeneration, swimming speed was directly related to the body wave characteristics of beat frequency, wavelength and wave speed (Fig. 5; Regression analysis, df = 1, P < 0.01). Multiple regression indicates that swimming speed depends on tail beat frequency (P < 0.001), but not on regeneration percentage (P = 0.65) or its interaction with tail beat frequency (P = 0.88) (Table S1, Fig. S1). Wave amplitude did not have a significant effect on swimming speed (Fig. 5D; Regression analysis, df = 1, P > 0.05). The beat frequency of the control sham lampreys was low compared to transected lampreys swimming at a similar speed; therefore, the control lampreys were able to achieve higher swimming speeds at lower beat frequencies and wave speeds than the transected lampreys (Fig. 5A and C; Oliphint et al. 2010).

**Fig. 5.**
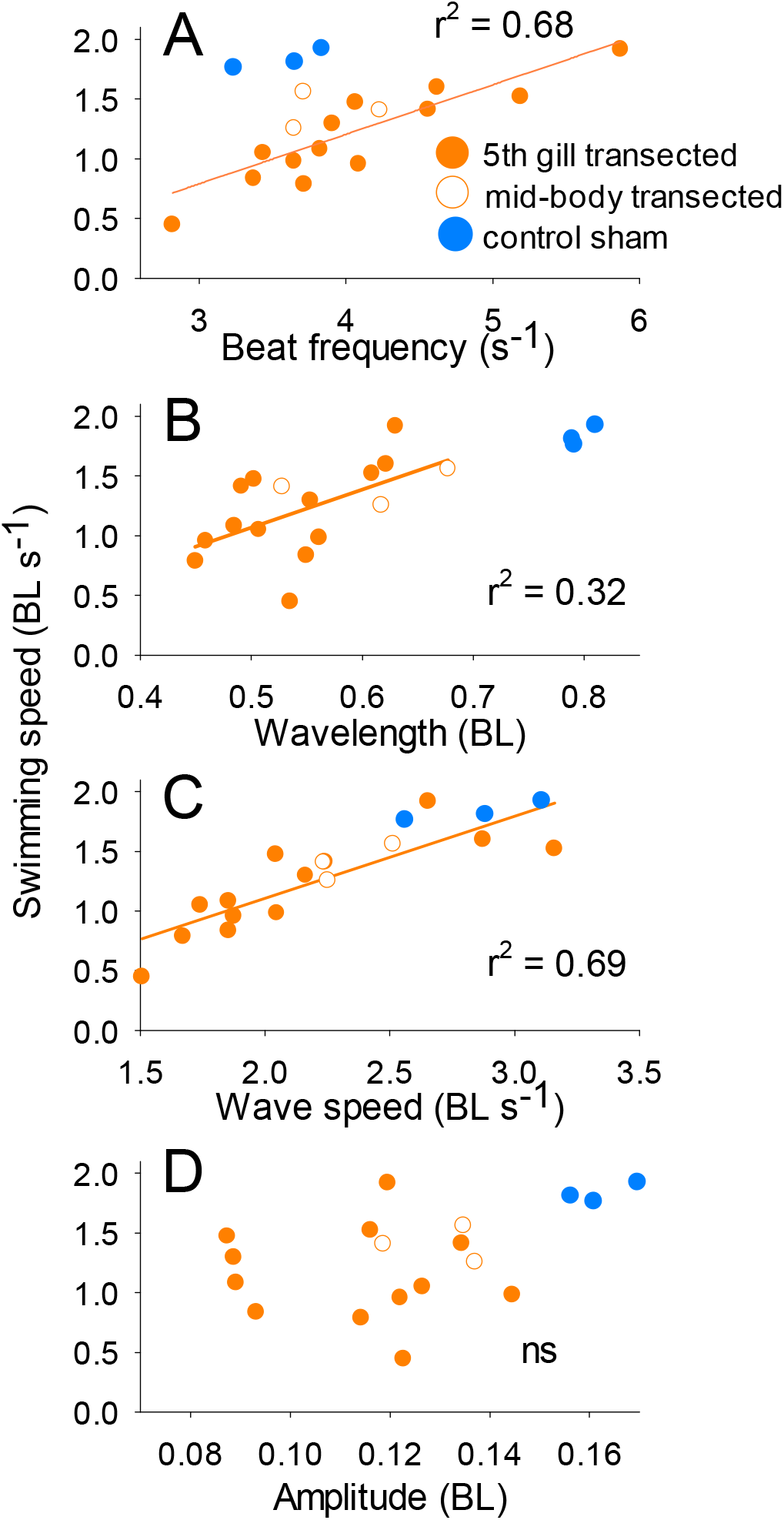
Body kinematic variables versus swimming speeds of different lampreys of the 5^th^ gill transected (filled orange circles), the mid-body transected (open circles) and control lampreys (filled blue circles). A) beat frequency, B) wavelength and C) wave speed were all positively related to swimming speed for the transected lampreys (Regression analysis, p < 0.01). Wave amplitude of the traveling body waves was not significantly related to swimming speed (Regression analysis, p > 0.1).

### Kinematic indicators of swimming efficiency

In order to examine how the differences in kinematics and performance may translate into efficiency, we calculated Strouhal number and stride length, indices that can be used as indicators of efficiency (Fig. 6). The Strouhal numbers (St) of the control lampreys (and one mid-body lamprey) fell within the range (St = 0.25 − 0.35) that has been shown to provide the maximum propulsive efficiency (Fig. 6A; Taylor et al. 2003, Eloy 2012) and were significantly lower than the Strouhal of the 5^th^ gill transected lampreys (Fig.6B; Holm-Sidak post-hoc comparison, p < 0.05). However, the controls did not significantly differ from the mid-body transected lampreys (Holm-Sidak post-hoc comparison, p > 0.05). The control lampreys also swam further with each tail beat (Fig. 6C; One-way ANOVA, F = 39.8, p < 0.001). Therefore, both of these indices suggest that even when transected lampreys swim as fast as controls, they do not swim as efficiently.

**Fig. 6.**
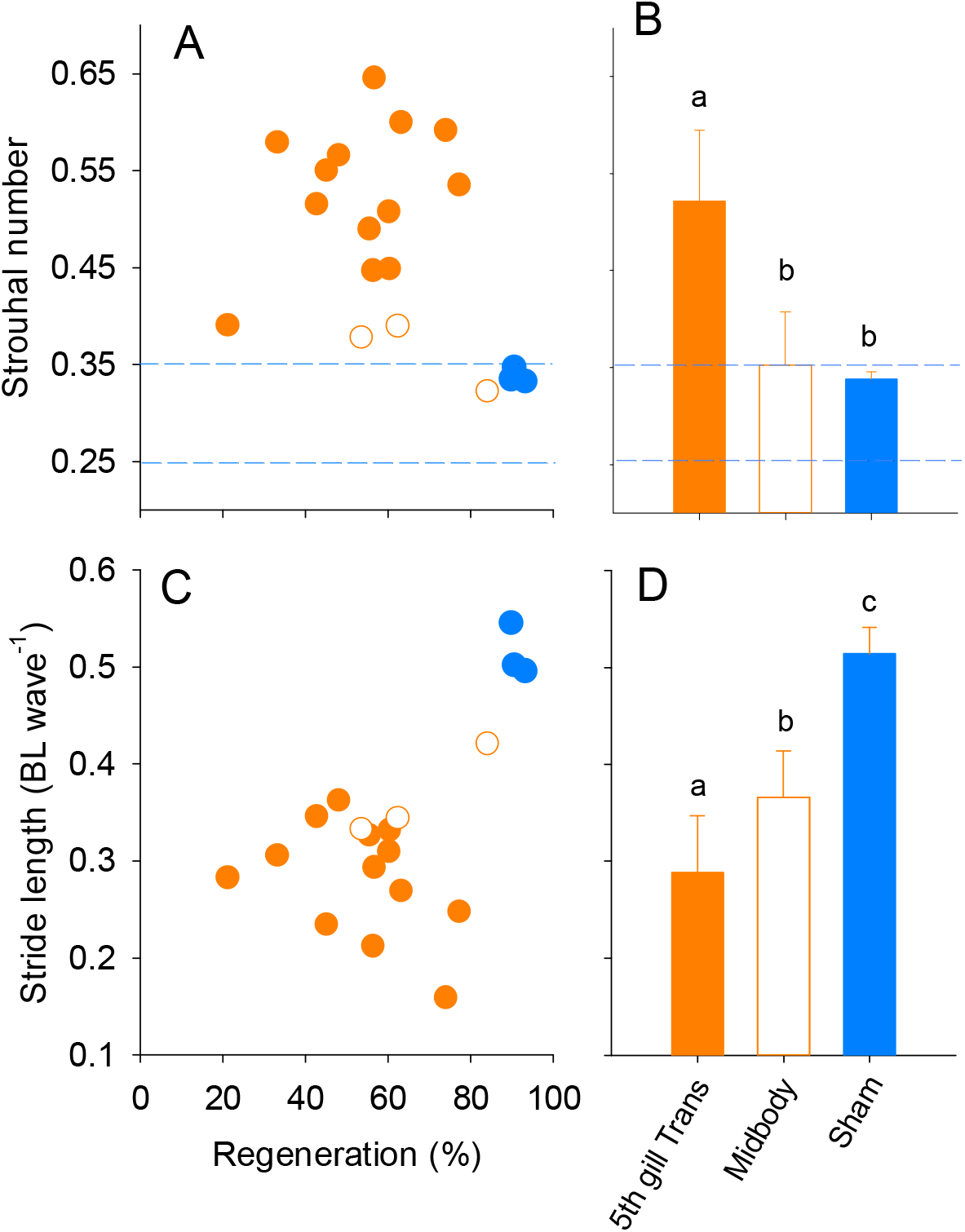
Effects of percent regeneration on the Strouhal number and stride length (BL traveled per wave) for the transected and control lampreys. A) Strouhal number of lamprey with different levels of regenerated spinal cord. Dotted blue lines highlight region where studies have shown animals and flapping foils to have the highest propulsive efficiency. The 5^th^ gill lampreys fell outside the optimal range of Strouhal while the control and the one mid-body lamprey fall within the optimal range. B) The 5^th^ gill (filled orange circles) transected lampreys had significantly higher Strouhal numbers than the mid-body (open circles) transected and control lampreys (Holm-Sidak post-hoc comparison, p < 0.05). C) Stride length of lampreys with different levels of regenerated spinal cord. D) Comparison of the stride lengths among treatments, letters designate significantly different groups (Holm-Sidak post-hoc comparison, p < 0.05).

### Comparison between 5^th^ gill and mid-body transected lampreys

The 5^th^ gill transected lampreys and the mid-body transected lampreys did not differ in any of measured performance or kinematic parameters (Figs. 1, 4, 5; Holm-Sidak post-hoc comparison, p > 0.05). However, the mid-body transected lampreys had significantly lower Strouhal numbers and stride lengths, compared to controls (Fig. 6D; Holm-Sidak post-hoc comparison, p > 0.05).

## Discussion

One of the primary goals of this study was to examine how swimming performance was related to degree of axon regeneration in lampreys recovering from spinal cord transection. We hypothesized that a larger fraction of axons regenerated would lead to more complete activation of the spinal locomotor circuits below the lesion. Based on this hypothesis, we predicted that animals with a greater fraction of regenerated axons would swim faster and more efficiently than those with fewer regenerated axons. But that is not what we observed. Swimming speed was not related to the percent axon regeneration of lampreys recovered from spinal cord transection (Fig. 2). However, individuals swam at highly variable speeds, whereby, most individuals could swim both rapidly and slowly (Supplemental fig. 1) and this variability may have obscured our ability to see a statistical relationship between swimming speed and axon regeneration. Basically, all the individuals, which had recovered for 10.5 weeks, had the ability to swim at variable speeds and modulated their swimming speeds by changing their wave frequency and shape (Fig. 5). Despite being able to swim, and, at times, swim relatively fast, transected individuals did not produce body waves as large as the control lampreys (having significantly lower wavelengths and amplitudes). As a result, in order to swim, fast transected individuals had to produce body waves very rapidly – i.e. high wave frequency – much higher frequency than control animals required to swim at the same speed. This suggests that the swimming efficiency, as indicated by Strouhal number and stride length was lower in the transected lampreys (both 5^th^ gill and mid-body) compared to control.

There appears to be a minimal recovery threshold required for lampreys to be capable of swimming, but there also seems to be a limit to how much they can recover. All the transected lampreys in this study, regardless of percent axon regeneration, which ranged from 33-84%, had a similar relationship between their wave frequency and swimming speed (i.e., similar stride length (Fig. 6B)). In fact, the wave kinematics and swimming performance of the 10.5 week recovered lamprey in this study were not much different than the 2 week recovered mid-body transected lamprey reported in Gemmell et al. (2016). It has been shown that spinal cord transected lamprey recover some locomotor function at 2 weeks, albeit with aberrant movements and locomotor activity and appear to increase their locomotor activity after that (Davis et al., 1993; McClellan, 1994). By 8 weeks recovered, transected lamprey have near normal locomotor movement and muscle activity patterns (McClellan, 1994). However, others have shown that even after 10 weeks, recovered lampreys need to use higher wave frequencies than control lamprey to reach similar swimming speeds (Oliphint et al., 2010). Likewise, we found that the recovered, transected lampreys in this study also swam significantly shorter distances per tail beat than the control lampreys. This suggests that recovered, transected lamprey are not capable of coordinating the kinematics necessary to generate swimming thrust as efficiently as non-transected lampreys.

Why are control lampreys able to swim better than transected lampreys? While all the lampreys in this study generated body waves that travel head to tail produced by waves of muscle activation on alternating sides of their body (McClellan et al., 2016; Williams, 1989), the shape and kinematics of these waves differ considerably between transected and control lampreys (Fig. 3). The body waves of the control lampreys are larger (longer wavelength and higher amplitude) and they develop more gradually resulting in amplitude increasing as each wave travels along the body (Fig. 4). The wave amplitudes of the transected lampreys did not increase gradually as the waves travelled along their bodies (Fig. 4). The gradual build-up of the wave amplitude has been shown to be essential for efficiently building and steering vortices for thrust generation (Gemmell et al., 2016). A comparison of the hydrodynamics generated by transected versus non-transected lampreys showed that the increase in wave amplitude gradually built up vorticity adjacent to the wave. The gradual build-up of vorticity lead to the non-transected lampreys generating suction thrust consistently along most of the body (Gemmell et al., 2015; Gemmell et al., 2016). In contrast, the body waves of the transected lampreys did not increase in amplitude or build vorticity along the body and thrust was inconsistent and primarily generated at the tail by positive pressure fields (Gemmell et al., 2016). Consequently, the non-transected lampreys get more thrust out of each body wave more efficiently (Fig. 6).

We speculate that transected animals, while they are able to produce muscle activity, are not able to produce as forceful contractions as control animals. Lower muscle forces would result in lower amplitude body waves, as we observed (Fig. 2E). Similarly, computational work has suggested that, when muscle forces are low compared to fluid forces, the body wavelength shortens (Tytell et al., 2010). If the wavelength of neural activity is similar in control and transected animals (as observed in vitro by McClellan, 1990), then the shorter mechanical body wavelength we observed would result in muscle activation earlier in the tail beat cycle relative to muscle shortening, and thus more eccentric activity, particularly toward the tail. Such eccentric muscle activity does not produce propulsive power, but instead may stiffen the caudal region to more effectively transmit muscle force from the anterior body to the fluid (Blight, 1977; Tytell et al., 2010). However, if the anterior body is not producing force effectively, as seems to be occurring in transected animals, the body stiffening may not be useful and may instead reduce the total power produced, decreasing swimming efficiency.

That lampreys can regain swimming behaviors post-recovery, despite incomplete axon regeneration, implies that other compensatory mechanisms are in play to restore locomotor behavior. In addition to RS axon regeneration, regeneration of other neuron types, as well as altered synaptic properties, has been observed within the lamprey spinal cord post-injury, which contribute to locomotor recovery (Becker and Parker, 2019; Cooke and Parker, 2009). Thus, the regenerated lamprey spinal cord is likely a “new” locomotor network (Parker, 2017).

In conclusion, just as there appears to be more than one way to “skin a cat” there appears to be more than one way for lampreys to swim. Recovered, transected lampreys clearly have the ability to swim and swim at high speeds. However, they have to produce many small body waves to achieve high swimming velocities which control lampreys achieve using less frequent, larger waves. The differences in wave kinematics rely on different thrust mechanisms (Gemmell et al., 2016) and ultimately result in different swimming efficiencies.

